# Multi-marker DNA metabarcoding detects suites of environmental gradients from an urban harbour

**DOI:** 10.1101/2022.04.17.488590

**Authors:** Chloe V. Robinson, Teresita M. Porter, Katie M. McGee, Megan McCusker, Michael T.G. Wright, Mehrdad Hajibabaei

**Affiliations:** Centre for Biodiversity Genomics and Department of Integrative Biology, University of Guelph, Guelph, Ontario, Canada N1G 2W1; Whales Initiative, Ocean Wise Conservation Association, Victoria, British Columbia, Canada V8V 4Z9; Environment and Climate Change Canada, Burlington, Ontario, Canada L7S 1A1

**Keywords:** Biomonitoring, metabarcoding, eDNA, bioindicators, COI mtDNA, 18S rRNA

## Abstract

There is increasing need for biodiversity monitoring, especially in places where potential anthropogenic disturbance may significantly impact ecosystem health. We employed a combination of traditional morphological and bulk macroinvertebrate metabarcoding analyses to benthic samples collected from Toronto Harbour (Ontario, Canada) to compare taxonomic and functional diversity of macroinvertebrates and their responses to environmental gradients. At the species rank, sites assessed using COI metabarcoding showed more variation than sites assessed using morphological methods. Depending on the assessment method, we detected gradients in magnesium (morphological taxa), ammonia (morphological taxa, COI sequence variants), pH (18S sequence variants) as well as gradients in contaminants such as metals (COI & 18S sequence variants) and organochlorines (COI sequence variants). Observed responses to contaminants such as aromatic hydrocarbons and metals align with known patchy distributions in harbour sediments. We determined that the morphological approach may limit the detection of macroinvertebrate responses to lake environmental conditions due to the effort needed to obtain fine level taxonomic assignments necessary to investigate responses. DNA metabarcoding, however, need not be limited to macroinvertebrates, can be automated, and taxonomic assignments are associated with a certain level of accuracy from sequence variants to named taxonomic groups. The capacity to detect change using a scalable approach such as metabarcoding is critical for addressing challenges associated with biodiversity monitoring and ecological investigations.

## Introduction

Ecosystem degradation is one of the leading causes of biodiversity decline in aquatic realms ^1^. Freshwater degradation can be observed as physical changes to habitat morphology, hydrological alterations and changes to biogeochemistry of water and sediment ^2,3^. Lakes in particular are more susceptible to the consequences of shoreline developments and loading of nutrients, including phosphorous and nitrogen, than other freshwater habitats ^4,5^. These lake stressors often cause a reduction in littoral habitats for submerged macrophytes, changes to sediment composition, and increased levels of eutrophication ^4–7^, resulting in loss of lake biodiversity ^2^. Restoring degraded lake ecosystems typically involves changing the ecological fate of the system towards an ecologically-sound status, where function is retained ^1^. To achieve this, stressors need to be first identified and then managed via intervention and ecosystem management techniques to enable recovery and restoration ^1,5,8^.

Toronto Harbour is located on the north shore of Lake Ontario, directly south of the City of Toronto and receives water from one of the most highly urbanized and industrialized areas in the Great Lakes. Contaminants from stormwater runoff, spills, and chemical input to sewers from industries and residences have contributed to severely degraded water and environmental health ^9–12^. In 1994, eight beneficial uses in Toronto Harbour were identified as impaired (beneficial use impairments; BUIs ^12,13^. In 1987, Toronto Harbour was designated as an Area of Concern (AOC) ^13^ and since then, there has been an increase in biological monitoring in the harbour ^9,14,15^.

As of 2016, results of Remedial Action Plan (RAP) activities resulted in re-classification of two of the eight identified BUIs (‘Not Impaired’), however “Loss of Fish and Wildlife Habitat’ remains impaired ^13,16^. Over the last 30 years, water quality, sediment quality and the quantity and condition of terrestrial and aquatic habitats have improved considerably, resulting in the preparation to delist Toronto Harbour as an AOC ^13^. In Toronto Harbour, the RAP investigates trends in benthic macroinvertebrate diversity by equating a higher diversity to a healthier system ^16^, with diversity metrics derived solely from morphological analyses ^17^. The challenges of morphological approaches compared to a DNA-based identification approach for monitoring macroinvertebrates have been highlighted previously (e.g. low taxonomic resolution and high processing costs) ^18–23^.

DNA extracted from bulk benthos samples has been transformative for routine benthic macroinvertebrate biomonitoring ^24^ and for determining gradients in freshwater environmental conditions ^18,25–28^. When taxonomic lists and corresponding the Hilsenhoff Biotic Index (HBI) tolerance values are paired with ecological information, such as functional feeding guilds (FFG), metabarcoding analysis can provide in-depth understanding of ecological function and responses of macroinvertebrates to different environmental conditions ^19,28,29,30,31^.

In this study, we hypothesized that the addition of eDNA metabarcoding will provide a more taxonomically comprehensive biodiversity measure for focal bioindicator macroinvertebrates, which will be reflected in finer-scale environmental assessment. By comparing diversity and functional macroinvertebrate metrics using eDNA metabarcoding and morphological assessment methods we assess the responses of these metrics across environmental gradients. Through this investigation, we aim to determine the most effective approach for long-term monitoring of benthic communities in Toronto Harbour and similar systems.

## Results

A map showing sampling stations in Toronto Harbour and associated gradients in water physical-chemical and sediment contaminants is shown in Figure 1 (Table S1). From these samples, a total of 18,790,832 x 2 paired-end sequence reads were generated for 75 samples and 16 controls. After read pairing, trimming, and denoising (sequence error, chimera, and pseudogene removal for COI) we generated a total of 9,329 exact sequence variants (ESVs) (Table S2).

**Figure 1.**
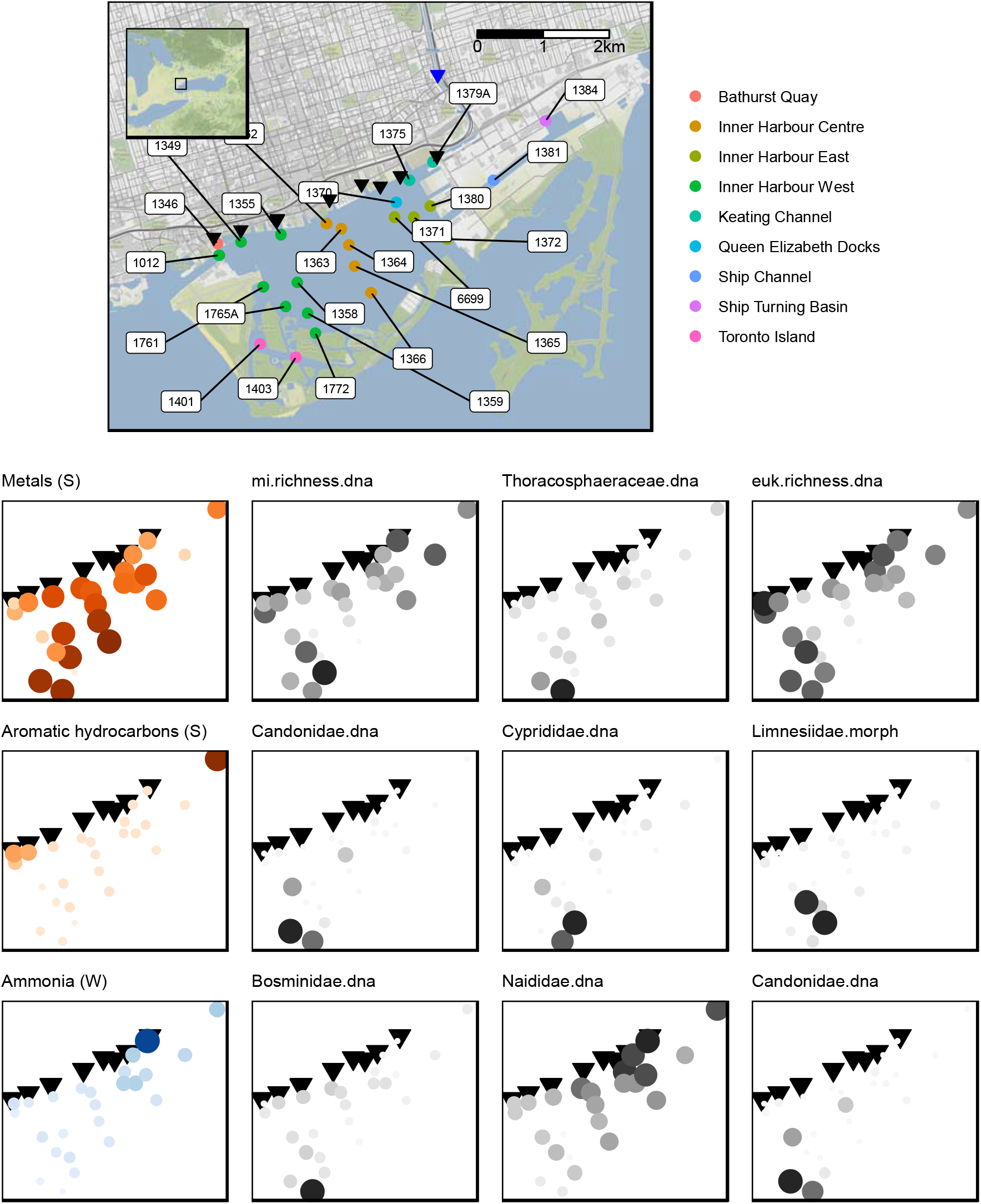
Toronto Harbour and associated abiotic and biotic gradients. Combined sewage outflows (black triangles) and the Don River (blue triangle) flows into the Toronto Harbour (Ontario, Canada). Sampling stations are shown in the legend. Inset map (top left) shows map area (black box) within respect to the Great Lakes region. Scale bar is in kilometers. Three of the most influential environmental variables are shown along with diversity metrics with the largest responses. Abbreviations: water physical-chemical features (W), sediment contaminants (S), macroinvertebrate (mi), eukaryote (euk). Map tiles by Stamen Design, under CC BY 3.0. Data by OpenStreetMap, under ODbL.

Overall, we found that taxonomic assignment resolution was finer using COI metabarcoding compared with other methods (Figure S1a). Sampling effort assessed using rarefaction found that sequencing depth was appropriate for the metabarcoding samples, and all curves reached saturation, however for samples identified by morphology, curves continue to rise indicating that further sampling would have identified additional families (Figure S1b).

### Macroinvertebrate diversity metrics measured using morphology or metabarcoding

Biodiversity was compared between morphological and metabarcoding approaches (Figure 2a). Macroinvertebrate species richness was higher using COI metabarcoding (median 15 species per station) compared with morphological methods (9 species per station) (Wilcox test, p-value > 0.00027) (Figure 2a). 18S metabarcoding detected an even higher diversity of genera (82 genera per station).

**Figure 2.**
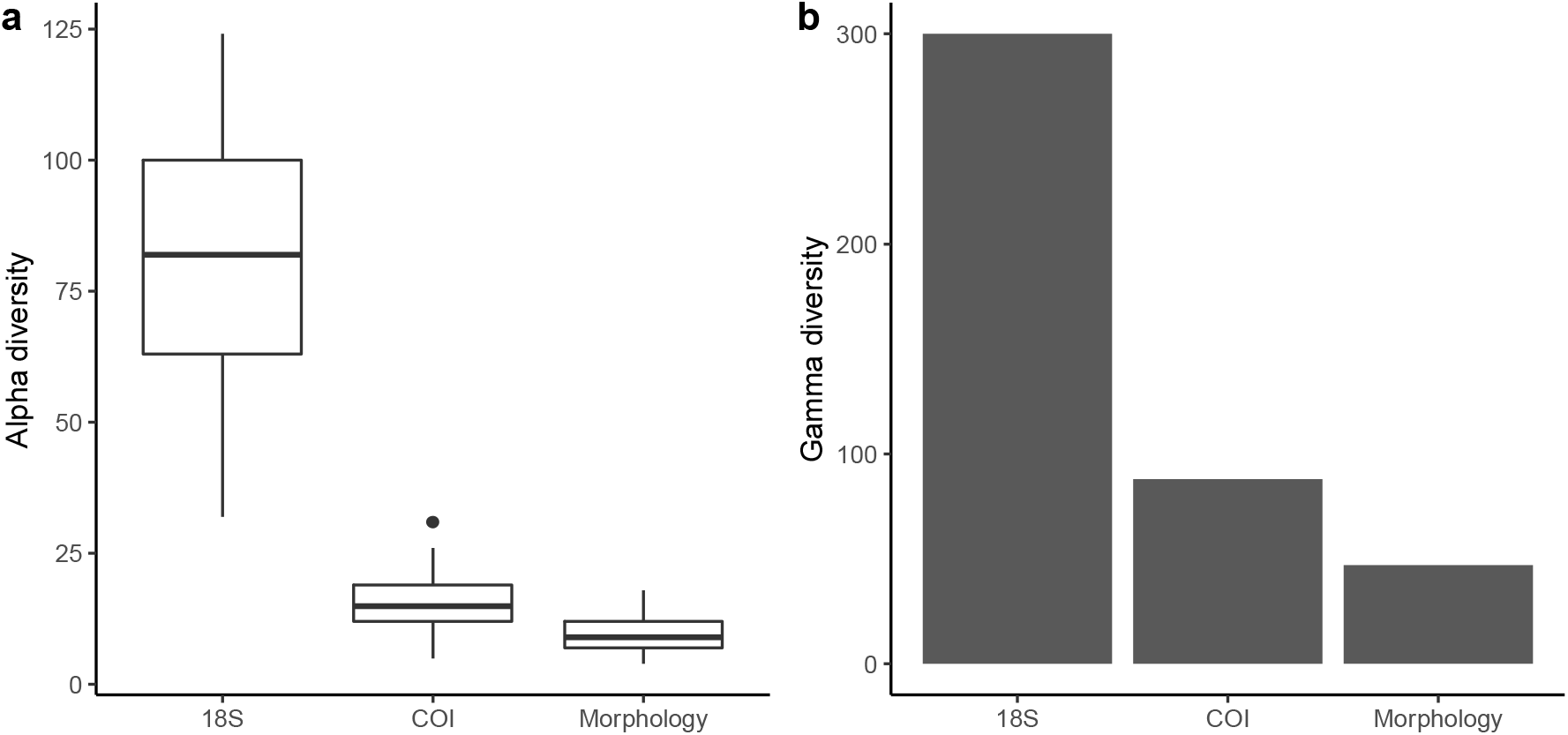
Alpha and gamma diversity detected is higher using DNA metabarcoding compared with morphological methods. In a) alpha diversity is compared across methods showing genus richness for (non-macroinvertebrate) eukaryotes sampled using 18S metabarcoding and species richness for macroinvertebrate sampling using COI and morphological sampling. In b) gamma diversity is compared showing the total number of 18S eukaryote genera and total number of species sampled using COI and morphological methods in Toronto Harbour.

The variance between the macroinvertebrate species detected using COI and morphological methods was similar for many stations, however, COI metabarcoding detected divergent communities at several stations (Figure 3a). The first two axes of the PCA plot explains 42.8% of the variance in these communities. These differences were correlated with the relative read abundance of *Limnodrilus hoffmeisteri, Tubifex tubifex*, and *Potamothrix vejdovskyi*. When we compared community composition using Bray Curtis dissimilarities, most locations appeared similar to each other when using morphological taxa (stress=0.09, R^2^=0,98) and ammonia (r = 0.31, p = 0.035) and magnesium (r = 0.35, p = 0.026) help explain the variance of these communities (Figure 3b). When we compared communities using COI sequence variants, locations showed more separation with Toronto Island and Bathurst Quay locations clustering separately from the rest (stress = 0.14, R^2^ = 0.86) and ammonia (r = 0.51, p = 0.002), metals (r = 0.38, p = 0.005), and organochlorines (r = 0.38, p = 0.005) help explain the variance of these communities (Figure 3c). When we compared communities using 18S sequence variants, most locations appeared similar, but Bathurst Quay and Keating Channel locations clustered separately (stress = 0.06, R2 = 0.99) and pH (r = 0.43, p = 0.004) and metals (r = 0.40, p = 0.005) help explain the variance of these communities (Figure 3d).

**Figure 3.**
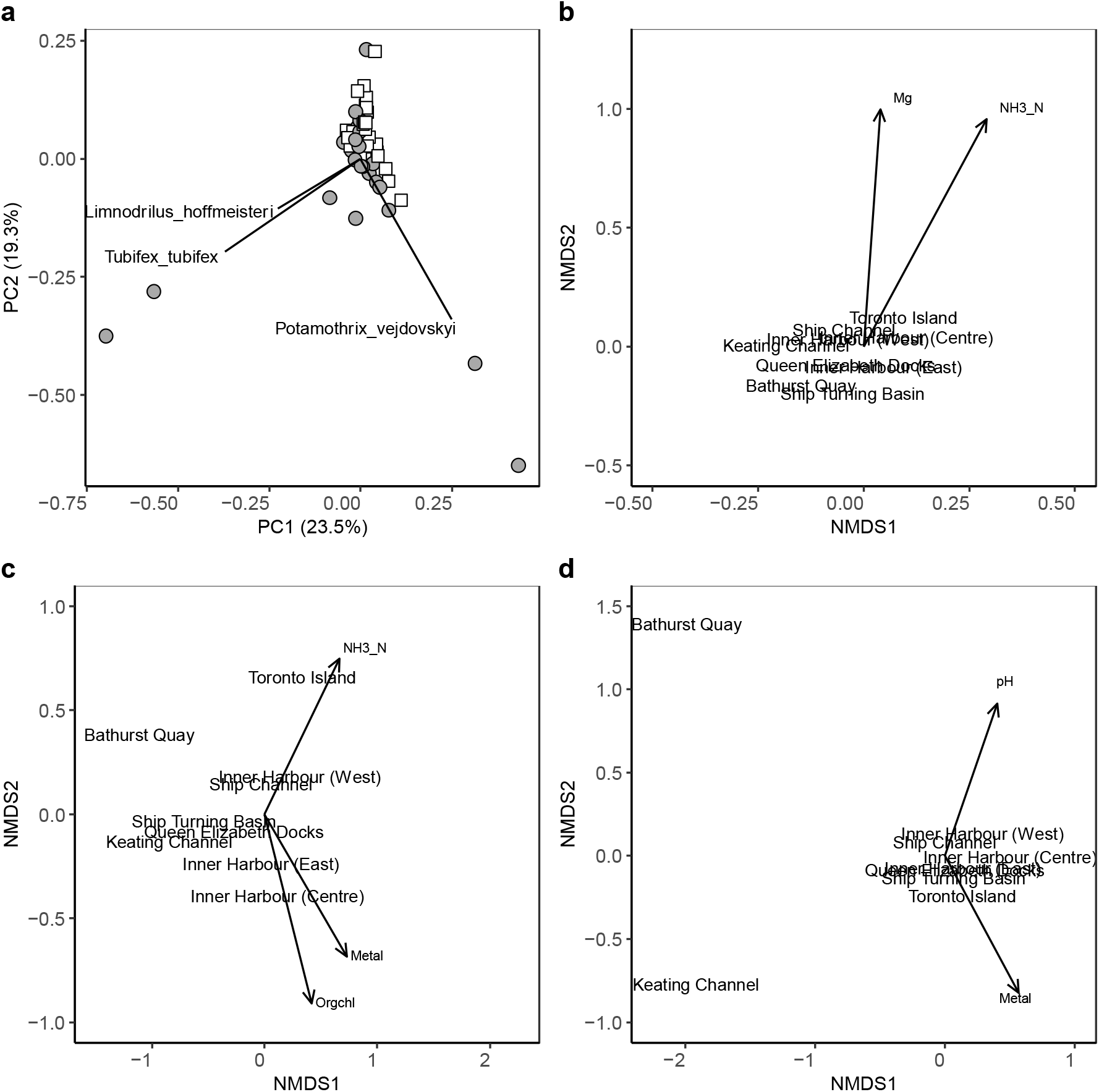
DNA metabarcoding helps distinguish between sampling locations in Toronto Harbour. Communities are compared using a) principal components analysis (PCA) at the species rank for COI and morphological sampling, b) non-metric multidimensional scaling (NMDS) using morphological taxa identified to a species rank when possible, c) NMDS using macroinvertebrate COI sequence variants, and d) NMDS using eukaryote (non-macroinvertebrate) 18S sequence variants. For PCA, COI samples are shown in grey circles and morphological samples in white squares and the most strongly correlated species vectors were added. For NMDS plots, centroids for each location are plotted for each sampling method using Bray Curtis dissimilarities, and environmental vectors were added if correlations were at least 0.30 with a p-value < 0.05.

We also compared the top 5 most abundant (non-macroinvertebrate) eukaryote families using 18S metabarcoding with the top 5 most abundant families detected using both morphological methods and COI metabarcoding (Figure 4a). Although we compare the relative abundance of individuals sampled using morphological methods with the relative read abundance of sequence variants, it’s important to acknowledge that read abundances reflect the specificity of our primers, specimen size, as well as abundance in samples. In terms of detections, COI metabarcoding detects some but not all the macroinvertebrate families detected using morphology. The median relative abundance of Naididae, the most abundant group using either method, was higher using morphological methods (90.6%) compared to COI metabarcoding (31.8%) (Wilcox test, p-value = 3.6e-10). The median relative abundance of Chironomidae was higher using COI (9.2%) compared to morphological methods (4.6%) (Wilcox test, p-value = 0.04). Overall, macroinvertebrate communities at the family rank were positively correlated (Pearson, 70.1%,p-value ~ 0, 95% confidence interval 61% - 78%).

**Figure 4.**
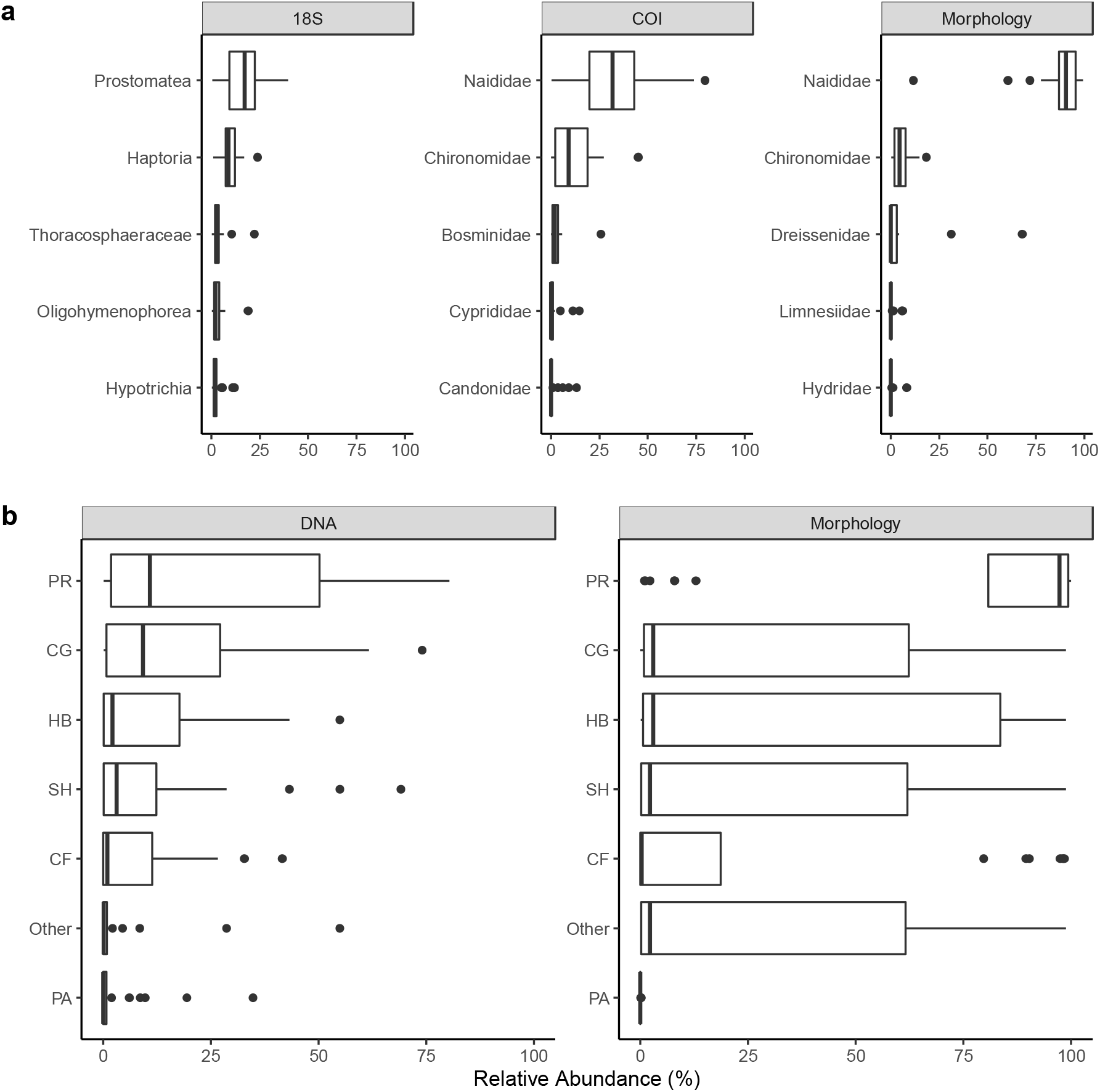
Comparison of major taxonomic and functional groups detected across sampling methods. We compare a) the 5 most abundant families of eukaryotes (non-vertebrate 18S) and macroinvertebrates (COI and morphology) and b) macroinvertebrate functional feeding guilds. Relative read abundance is presented for DNA-derived data and relative abundance of individuals is presented for morphology-derived data. Abbreviations: predators (PR), collector-gatherer (CG), herbivore (HB), shredders (SH), collector-filterers (CF), parasites (PA).

Overall, multi-marker metabarcoding recovers a greater diversity of taxa than morphological methods even when only considering macroinvertebrate taxa in the phyla Arthropoda, Annelida, Mollusca, Cnidaria, and Platyhelminthes for a fair comparison with morphological methods (Table S3). Overall, we detected 47 and 88 unique macroinvertebrate species (using morphological methods versus COI metabarcoding), 77 and 79 genera, and 30 and 61 families in total across all sampled stations. When considering other (non-macroinvertebrate) eukaryotes detected using 18S metabarcoding, we detected a further 271 genera and 111 families from 36 eukaryote phyla with the most diverse phyla detected from the Ciliophora (105 taxa), Cercozoa (64), Ascomycota (Fungi, 61), Basidiomycota (Fungi, 52 taxa), and the Chytridiomycota (42 taxa).

### Macroinvertebrate functional metrics using metabarcoding and morphological methods

Functional feeding groups, a classification system based on how macroinvertebrates acquire food, allowed us to assess whether ecological function varies independently of taxonomic shifts. At each station, the proportion of macroinvertebrate reads (COI metabarcoding) or individuals (morphological) assigned to a family with a particular functional feeding guild(s) was assessed (Figure 4b). The relative abundance of FFGs between methods were significantly different for parasites (Wilcox test, = 0. 3.8e-05), predators (p.adj = 0. 1.8e-05), and ‘other’ (p.adj = 0.014). The median relative abundance of parasites was ~0% using morphological methods and 0.03% using COI metabarcoding. The median relative abundance of predators was 97% using morphological methods and 11% using COI metabarcoding. The mean relative abundance of ‘other’ was 2% using morphological methods and 0.2% using COI metabarcoding. Functional communities detected using both methods were not found to be correlated (Pearson, 0.08, p-value = 0.31, 95% confidence interval - 0.07 - 0.22), though this may be due to our difficulty assigning function to macroinvertebrate families detected using COI metabarcoding methods. The three most abundant predator families using COI metabarcoding was Naididae (422 sequence variants, 284,851 reads), Chironomidae (81 sequence variants, 96,261 reads), and Bosminidae (34 sequence variants, 23,495 reads); and using morphological methods was Naididae (30 taxa, 30,163 individuals), Chironomidae (41 taxa, 322 individuals), Dreissenidae (3 taxa, 2241 individuals).

We used hierarchical partitioning to identify environmental variables that explain the variance in diversity and functional metrics (Table S5, Table S6). Sediment contaminants explain more variation in diversity metrics and water physical-chemical features tend to explain more variation in functional metrics (Figure 5). The most influential environmental parameters are metals, aromatic hydrocarbons, and ammonia (Figure 1). Metals make significant independent contributions explaining 62% of the variation in macroinvertebrate species richness (COI), 55% of the variation in the relative abundance of Thoracosphaeraceae (18S), and 41% of the variation in eukaryote genus richness (18S). Aromatic hydrocarbons make significant independent contributions explaining 63% of the variation in Candonidae (COI), 57% of the variation in Cyprididae (COI), 45% of the variation in Limnesiidae (morphological), and 41% of the variation in Naididae (morphology). Ammonia makes significant independent contributions explaining 56% of the variation in Bosminidae (COI), 56% of the variation in Naididae (COI), 50% of the variation in Candonidae (COI), 49% of the variation in Thoracosphaeraceae (18S), and 47% of the variation in Hydridae (morphological). Temperature makes significant independent contributions explaining 62% of the variation in macroinvertebrate richness (COI) and 50% of the variation in Haptoria (18S). Ammonia makes significant independent contributions explaining 57% of the variation in parasites (morphology), 30% of the variation in shredders (morphology), 29% of the variation in collector-gatherers (morphology). Organochlorines also make significant independent contributions explaining 59% of the variation in parasites (COI) and 46% of the variation in collector-filterers (COI).

**Figure 5.**
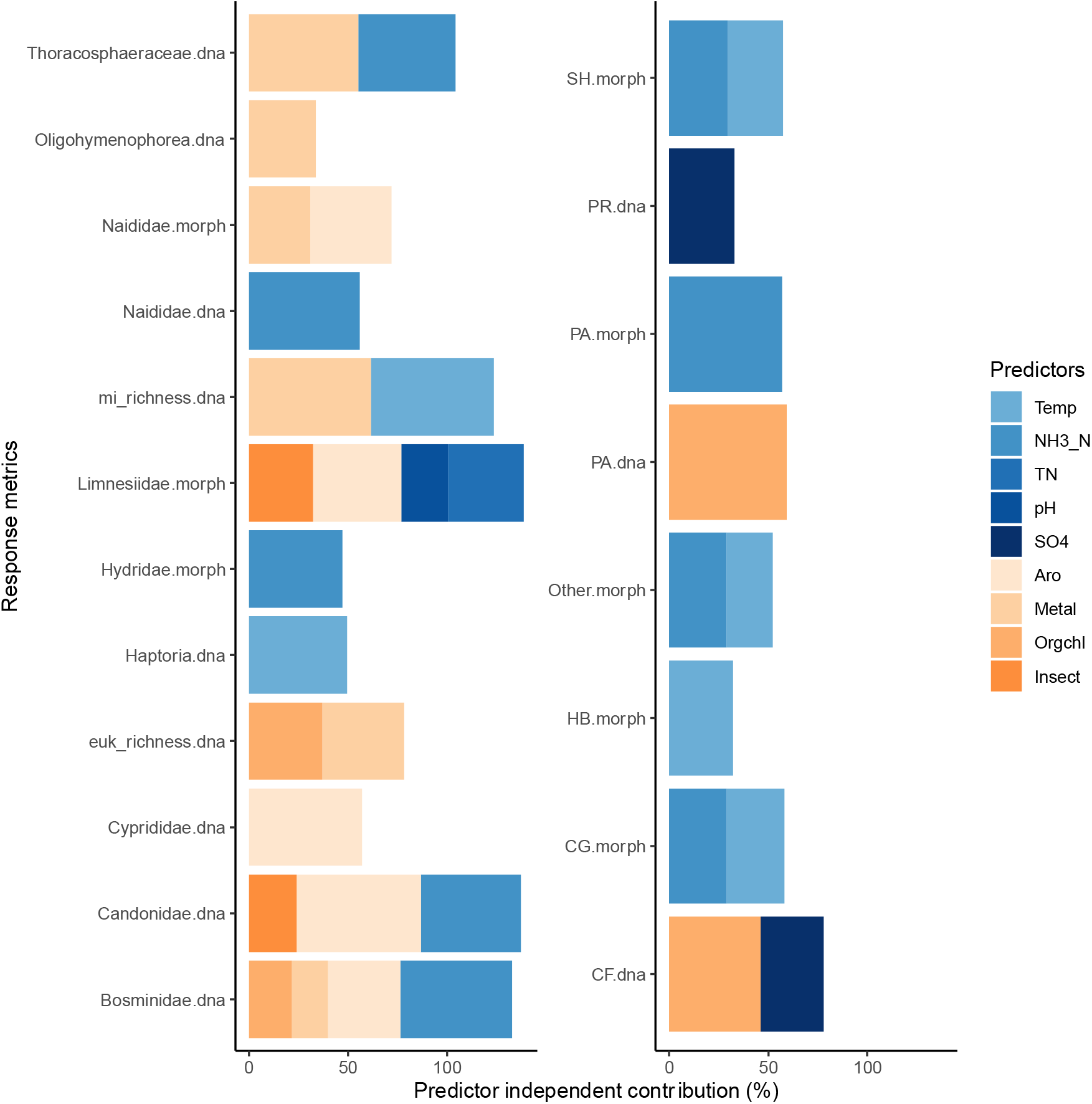
Sediment contaminants explain more variation in diversity metrics and water physical chemical features explain more variation in functional metrics. Predictors that make significant (p-value < 0.05) independent contributions explaining variance in diversity metric response variables (left panel) and functional metric response variables (right panel). Abbreviations: DNA metabarcoding (dna), macroinvertebrate (mi), eukaryote (euk), conventional morphology (morph), shredders (SH), predators (PR), parasites (PA), herbivores (HB), collector-gatherers (CG), collector-filterers (CF), temperature (Temp), ammonia (NH3_N), total nitrogen (TN), sulphate (SO4), aromatic hydrocarbons (Aro), metals (Metal), organochlorines (Orgchl), insecticides (Insect).

## Discussion

The process of monitoring benthic macroinvertebrate communities has been the focus of freshwater health assessments across the world. Morphological-based approaches, which were once the only method of determining richness, are rapidly being complemented with DNA-based methodologies such as metabarcoding, greatly increasing the taxonomic information obtained from benthic samples ^21,21,32–34^. In this study, we have demonstrated the benefits of employing eDNA metabarcoding to lake-based macroinvertebrate biodiversity assessments.

Previously, morphological-based biomonitoring in Toronto Harbour focused primarily on overall richness as the determining factor of lake health ^15,16,35^ with limited investigation into the composition of taxa. While this can be informative as to the overall quantity of macroinvertebrate taxa present in the lake, it misses details on the proportions of bioindicator groups in relation to other taxa. Moreover, species diversity often decreases with degradation of aquatic environments, with more tolerant species dominating species compositions and composition of macroinvertebrates ^36^. Therefore, the concept of higher total richness translating to improvements in water quality and lake health are misleading, as measures of community composition, structure and diversity are also required to understand changes in environmental gradients ^36^. COI metabarcoding also provides finer level taxonomic assignments compared with morphological methods as has been shown in a previous study for fishes ^37^.

In addition, the confidence levels for morphological macroinvertebrate identification are scarcely reported and classifications can vary between taxonomists ^18,38^, meaning likelihood of misidentified or missed taxa is not taken into consideration ^21^. Both DNA metabarcoding (i.e., finer resolution taxonomic assignments, ability to detect microscopic taxa) and morphological methods (i.e., absolute abundance counts) provide complementary information that can contribute and support macroinvertebrate community information for further ecological analyses. In addition to the typical benthic macroinvertebrates identified using morphology, DNA metabarcoding enabled accurate detection of additional phyla, using more inclusive markers as used in this study. In this study, the detection of unique taxa via 18S sequencing enabled us to clearly distinguish stations, highlighting that non-macroinvertebrate groups can also provide insights towards understanding of spatial community dynamics in lake systems ^39^.

The 18S marker enabled us to detect fungal groups expected to be found from aquatic environments such as the Chytridiomycota, Blastocladiomycota, and Cryptomycota (Rozellomycota). We even detected members of the Archaeorhizomycota (12 sequence variants, 117 reads), a group of fungi with a global distribution largely known only from marker gene studies of soil with only two species in culture ^40,41^. The application of this marker also resulted in detection of several fungal taxa that we would expect to find from benthic samples such as known parasites of arthropods/insects: *Cordyceps, Coelomomyces*, Labulomycetales, and Cryptomycota.

Compared to DNA methods, morphological methods detected negligible amounts of aquatic bioindicator taxa, specifically Ephemeroptera (HBI= 0-2; excellent) and Trichoptera (HBI= 0-4; very good). COI metabarcoding detected Ephemeroptera and Trichoptera more often, but the relative abundance of these taxa were relatively low. Environmentally tolerant taxa (i.e. those with HBI score of 6-10), were detected more so with DNA metabarcoding methods compared to morphology. For example, species/genera including, *Limnodrilus hoffmeisteri* (HBI= 10; very poor), and *Tubifex tubifex* (HBI = 10; very poor), and *Hydra* (HBI = 5; poor) were detected exclusively by DNA methods, whereas *Potamothrix vejdovskyi* (HBI= 8; poor) was detected exclusively with morphological methods. Naididae as a family are highly tolerant to organic pollution, scoring between 6 and 10 on the HBI index ^42^. More families, genera and species from extreme ends of the tolerance index were detected using DNA metabarcoding, highlighting the ability of this method to detect presence of these important taxa.

Beyond taxonomic metrics, mapping functional diversity across lake environments can improve our understanding of lake ecosystem integrity ^19,43–45^. For example, the presence of diverse macroinvertebrate predators detected in Toronto Harbour may indicate that there are sufficient resources to support a stable multi-level food web ^43^. Overall, morphological methods performed very well with regards to their ability to identify a diverse array of functional groups, but it was challenging to use automated methods to annotate function to many of the less abundant but diverse macroinvertebrate families detected using COI metabarcoding. Despite metabarcoding methods facilitating genus- and species-level taxonomic assignments, databases of functional metrics for macroinvertebrates at this taxonomic level are still incomplete ^46^. The influence of different environmental variables (e.g. Metals and aromatic hydrocarbons in sediment as well as ammonia in lake water) on the relative abundance of functional feeding groups has implications on the trophic system of the lake as a whole. An increase or decrease in sediment and water pollutant loads will likely influence a shift in the taxa exhibiting different FFG guilds. Assigning FFG at more conservative taxonomic ranks (i.e., family) enables easy integration of both metabarcoding and morphological taxonomic lists whilst maximizing functional assignments.

By using a multi-marker approach, we can begin to close knowledge gaps regarding how water quality and loading of elements in sediment affect taxa at various trophic levels ^47,48^. In this study, metals were found to be one of the more influential sediment contaminants explaining variation in overall macroinvertebrate richness using COI metabarcoding as well as variation in (non-macroinvertebrate) eukaryote richness using 18S metabarcoding. In Toronto Harbour, metals such as cadmium, chromium, copper, lead, mercury, and zinc were found to have levels that exceeded the Canadian Federal Probable Effect Level. In this study, we have samples from 5 stations from inner harbour centre that represent a clear gradient of increasing metal concentrations, decreasing macroinvertebrate and eukaryote richness detected using COI and 18S metabarcoding. Ciliates, despite their ubiquity in freshwater systems, the key position they play in trophic webs (feed on bacteria, algae, other protists while also being consumed by other meiofauna) and known sensitivity to a range of water quality types (low to highly polluted), they are often a neglected in water quality assessments ^49^.

## Conclusion

DNA methods can readily be applied for both routine lake community biomonitoring and targeted monitoring for invasive, threatened, and/or exploited lake species, integrated with periodic morphological assessments to supplement routine DNA monitoring with abundance measures. Overall, for taxonomically comprehensive assessments of lake communities, it is imperative to apply robust high-throughput methods, such as DNA metabarcoding, to increase the resolution of biodiversity data and to understand species-specific responses to environmental gradients in the face of various perturbations and climate warming effects.

## Methods

### Study stations and sampling design

Sample stations in Toronto Harbour have been previously established by the Ministry of the Environment and Climate Change Resources in 1971, to track improvements in sediment contaminant concentrations and benthic invertebrate species composition and density ^50^ (Table S1; Figure S1).

In October 2018, water, sediment and benthos samples were collected from Toronto’s Inner Harbour over a three-day period, from a subset of 25 stations ^17^. At each station, environmental data was collected, followed by the collection of surface and overlying water, sediment and benthos samples in order. Samples were collected following the Canadian Aquatic Biomonitoring Network (CABIN) Open Water sampling protocols for collection of benthic samples ^51,52^.

Water samples were analyzed for major ions, nutrients, temperature, conductivity, pH and dissolved oxygen (see supporting information for technical details). Sediment samples were analyzed for trace metals, PCB aroclors, total PCBs and PCB congeners, chlorinated pesticides, chlorobenzenes, technical toxaphene (insecticide) and toxaphene congeners/parlars. See supporting information for technical details on water, sediment and benthos sampling and collection for both DNA and morphological identification.

### Morphology-based taxonomic identification of benthos

Invertebrates in the benthic community samples were sorted, identified to the family level, and counted by EcoAnalysts, Moscow, ID.

### DNA sample homogenization and DNA extraction of benthos samples

Samples were transferred from whirl-packs to 50mL conical tubes, using molecular biology grade water to rinse whirl packs to ensure the entire sample was transferred. Samples were centrifuged to collect sediment at the bottom, and excess water was removed. Approximately 0.3g was then directly subsampled into 2mL bead tubes included within the Qiagen PowerSoil kit. Samples were extracted according to manufacturer’s protocol eluting with 50uL Buffer C6. Each batch (~95 samples) included one negative extraction control where no tissue was included. The remaining mass was stored in the Falcon tubes at −20°C as a voucher.

### Library preparation and high-throughput sequencing

Two fragments within the standard COI DNA barcode region and one fragment within the 18S (eukaryote) region were amplified with the following previously optimized and validated primer sets: (LCO1490/230_R [~230 bp] called F230R, mICOIintF/jgHCO2198 [~313 bp] called ml-jg and Uni18SF/ Uni18SR [~600 bp] called Uni18S ^53–57^, using a two-step PCR amplification regime. The first PCR used COI and 18S specific primers and the second PCR involved Illumina-tailed primers. Uni18S PCR cycling conditions are as follows: 95°C for 3 minutes, followed by 35 cycles of 94°C for 30 seconds, 52°C for 30 seconds, 72°C for 90 seconds, and a final extension of 72°C for 8 minutes. COI PCR cycling conditions for both fragments were: 95°C for 5 minutes, followed by 35 cycles of 94°C for 40 seconds, 46°C for 1 minute, 72°C for 30 seconds, and a final extension of 72°C for 5 minutes. Amplification was visually confirmed through a 1.5% agarose gel electrophoresis. Amplicons were purified with a MinElute PCR purification kit (Qiagen), quantified with fluorometry using a QuantIT PicoGreen dsDNA assay kit (Invitrogen), and normalized to the same concentration prior to dual indexing with a Nextera Index Kit (Illumina). Indexed samples were pooled and purified through magnetic bead purification. The library was then quantified with the QuantIT PicoGreen dsDNA assay kit, and average fragment length was determined on an Agilent Bioanalyzer 2100 with a DNA 7500 chip. The library was then diluted to 4nM based on the concentration and average fragment length and sequenced on an Illumina MiSeq using a v3 chemistry kit (2 x 300 cycles). A 10% PhiX control spike-in was used to ensure sequence diversity.

### Bioinformatic processing

Illumina MiSeq paired-end reads were processed using the MetaWorks-1.0.0 pipeline available from https://github.com/terrimporter/MetaWorks ^58^. MetaWorks is an automated Snakemake ^59^ bioinformatic pipeline that runs in a conda ^60^ environment. Details of the sequence processing steps of METAWORKS is described in the Supplementary Material. Further analysis of leave-one-sequence-out testing with the RDP Classifier conducted with our COI and 18S reference sets allowed us to assess expected taxonomic assignment accuracy according to metabarcode length and taxonomic rank ^61–63^. Using a leave one sequence out approach during classifier validation, we determined that for a ~ 200 bp COI metabarcode, taxonomic assignments are ~ 90% correct at the species rank, 95%+ correct at the genus - family ranks using the appropriate bootstrap support cutoffs, and 99%+ correct at more inclusive ranks (e.g. order - kingdom) assuming the query taxa are present in the reference database ^61^. Using a ~ 200 bp 18S metabarcode, taxonomic assignments are ~ 80%+ correct at the genus - order ranks using the appropriate bootstrap support cutoffs and about 95%+ correct at more inclusive ranks (class - domain) assuming the query taxa are present in the reference database.

### Statistical analyses

All statistical analyses were conducted in RStudio v 1.1.456 running R v 3.5.1 ^64^ and plots were created with ggplot2 ^65^. Sequence variants recovered from field blanks comprised 2.7% of unique COI macroinvertebrate sequence variants and 5% of unique 18S eukaryote sequence variants and were removed from all subsequent analyses.

Resolution of taxonomic assignments were compared by recoding taxonomic assignments to: species=1, genus=2, family=3, etc. Sampling effort in metabarcoding and morphological samples were assessed using rarefaction. Relative abundance were compared by calculating the proportion or percentage of reads per taxon per station for metabarcoding data and by calculating the number of individuals per taxon per station for morphological methods. Prior to NMDS or PCA matrices were converted to proportions or standardized using a Hellinger transformation.

For a fair comparison of macroinvertebrate metrics using COI metabarcoding and morphological methods, we limited COI results to taxa in the phyla normally detected using morphological methods: Arthropoda, Annelida, Mollusca, Cnidaria, and Platyhelminthes. We compared several macroinvertebrate diversity metrics for morphology-based and metabarcoding methods such as richness at the species level for macroinvertebrate COI and morphological samples, richness at the genus level for (non-macroinvertebrate) eukaryote 18S samples and read abundance of the top 5 most abundant families per sampling method.

We used principal components analysis (PCA) ‘rda’ function in the vegan package to assess the variance between macroinvertebrate COI metabarcode and morphological samples ^66^. Only vectors for the most strongly correlated species were plotted for clarity. Also using vegan, we used non-metric multi-dimensional scaling (NMDS) analyses to visualize beta diversity based on Bray Curtis dissimilarities using the ‘metaMDS’ function using 3 dimensions and fitted correlated environmental variables using the ‘envfit’ function using 999 permutations if correlations were greater than 0.30 and the p-value < 0.05. Goodness of fit calculations and Shephard’s curve were calculated using the vegan ‘goodness’ and ‘stressplot’ functions.

Water physical-chemical features were measured from samples collected at the bottom of the harbour and contaminants were measured from sediment samples. For each set of measurements for organic contaminants/metals, values for each individual chemical/element was summed across each major contaminant group ^17^. We tested each predictor for normality using a Shapiro-Wilk Test using the ‘shapiro.test’ function in R. Skewness was checked using the ‘skewness’ function from the moments package ^67^. Each individual predictor variable was then transformed (square root, log10, 1/x) as needed to better meet normality assumptions in hierarchical partitioning using a Gaussian model. Predictors were standardized using z-scores and centered prior to analysis. We checked for collinearity among water and sediment variables using Pearson correlation coefficients with a cutoff of 0.70, using the ‘rcorr’ function from the Hmisc package (Figure S2)^68^.

For each family, we added primary functional feeding guild (FFG) annotations based on the EPA Freshwater Biological Traits Database ^69^. This system recognizes 7 feeding modes: collector-filterer (CF), collector-gatherer (CG), herbivore (scraper) (HB), parasite (PA), predator (piercer, engulfer) (PR), shredder (SH), and Other. Remaining missing family annotations were added using information compiled from the Taxa and Autecology Database for Freshwater Organisms available from freshwaterecology. info ^70^. For this database, preference was given to feeding type annotations by Moog ^71^. Any further missing annotations were then added by checking feeding habit annotations Since individuals were identified to the family level, species in a family can exhibit more than one feeding type and the presence of multiple feeding types per family were recorded when this was the case. We multiplied a family x station matrix containing read/individual counts by each feeding mode in a family x FFG matrix containing 1’s or 0’s indicating the presence of the feeding mode in a family. For each feeding mode, total read counts per station were recorded and combined into a single FFG x station matrix. Out of 61 families detected using metabarcoding, 23 (38%) were assigned feeding types. Out of 30 families detected using morphology, 24 (80%) were assigned feeding types. Further analyses using FFGs represent the proportion of all reads/individuals that could be both taxonomically assigned to family and functionally assigned a feeding mode.

We used hierarchical partitioning to regress each macroinvertebrate metric, individually, on the environmental predictors to identify which water physical-chemical features or sediment contaminants explain the most variance independently of the others. This was done using the ‘hier.part’ package in R ^73^.

Nutrient/major ion predictors measured from water collected at the bottom of the harbour were analyzed separately from sediment contaminants. Macroinvertebrate diversity metrics included macroinvertebrate species richness based on COI and morphological methods, genus richness based on (non-macroinvertebrate) eukaryote 18S metabarcoding, as well as the relative abundance of the top 5 families based on COI, morphological, and 18S. Functional metrics included the FFGs calculated for macroinvertebrate families using COI and morphological methods. All metrics were standardized using a Hellinger transformation using the ‘decostand’ function in vegan. Hierarchical partitioning significance was assessed using a randomization test, 1000 replicates, and calculating a goodness of fit measure based on log-likelihood.

## Acknowledgments

We would like to thank Adam Morden, Jonathan Paynter, Tina Mamone, and Jesse Baillargeon from ECCC for field support and members of the Hajibabaei lab for help processing samples. Funding for sampling and data collection for this project was provided by Environment and Climate Change Canada’s Great Lakes Action Plan in conjunction with the government of Canada through Genome Canada and Ontario Genomics.

## Author contributions

MM and MH designed the study, MM collected samples, MTGW processed and sequenced samples, CVR and TMP conducted bioinformatic and statistical analyses with assistance from KMM, CVR wrote the manuscript with help from all co-authors.

## Data availability

Raw sequences are available from the NCBI SRA PRJNA835155. The bioinformatic pipeline MetaWorks v1 is available from GitHub at https://github.com/terrimporter/MetaWorks/releases/tag/v1.0.0, the COI Classifier v4 and the 18S Classifier v4.1 we used are available from GitHub at https://github.com/terrimporter/CO1Classifier and https://github.com/terrimporter/18SClassifier respectively. The code used to generate figures, including infiles, are available from https://github.com/Hajibabaei-Lab/RobinsonEtAl2022_TorontoHarbour.

## Additional information

The authors declare that they have no known competing financial interests or personal relationships that could have appeared to influence the work reported in this paper.

## Notes

### Competing Interest Statement

The authors have declared no competing interest.

### Summary of Updates

Figures 1 and 5 revised; author affiliations updated; supplementary tables updated; 'Macroinvertebrate functional metrics using metabarcoding and morphological methods' section updated to clarify environmental contributions; discussion updated to reflect changes to 'Macroinvertebrate functional metrics using metabarcoding and morphological methods' section

## References

1. Breed, M. F. et al. The potential of genomics for restoring ecosystems and biodiversity. Nat. Rev. Genet. 20, 615–628 (2019).

2. Carpenter, S. R., Stanley, E. H. & Vander Zanden, M. J. State of the world’s freshwater ecosystems: Physical, chemical, and biological changes. Annu. Rev. Environ. Resour. 36, 75–99 (2011).

3. Geist, J. Integrative freshwater ecology and biodiversity conservation. Ecol. Indic. 11, 1507–1516 (2011).

4. Jeppesen, E., Søndergaard, M., Meerhoff, M., Lauridsen, T. L. & Jensen, J. P. Shallow lake restoration by nutrient loading reduction--some recent findings and challenges ahead. Hydrobiologia (2007).

5. Søndergaard, M. & Jeppesen, E. Anthropogenic impacts on lake and stream ecosystems, and approaches to restoration. J. Appl. Ecol. 44, 1089–1094 (2007).

6. Marburg, A. E., Turner, M. G. & Kratz, T. K. Natural and anthropogenic variation in coarse wood among and within lakes. J. Ecol. 94, 558–568 (2006).

7. Schindler, D. W. Recent advances in the understanding and management of eutrophication. Limnol. Oceanogr. 51, 356–363 (2006).

8. Lau, S. S. S. & Lane, S. N. Continuity and change in environmental systems: the case of shallow lake ecosystems. Prog. Phys. Geogr. Earth Environ. 25, 178–202 (2001).

9. Brinkhurst, R. O. Distribution and abundance of Tubificid (Oligochaeta) species in Toronto harbour, Lake Ontario. J. Fish. Res. Board Can. 27, 1961–1969 (1970).

10. Wood, L. W. & Chua, K. E. Glucose flux at the sediment-water interface of Toronto Harbour, Lake Ontario, with reference to pollution stress. Can. J. Microbiol. 19, 413–420 (1973).

11. Nriagu, J. O., Wong, H. K. T. & Snodgrass, W. J. Historical records of metal pollution in sediments of Toronto and Hamilton harbours. J. Gt. Lakes Res. (1983).

12. Toronto & Region Remedial Action Plan. Metro Toronto and Region Remedial Action Plan. (1989).

13. Dahmer, S. C., Matos, L. & Morley, A. Restoring Toronto’s waters: Progress toward delisting the Toronto and Region Area of Concern. Aquat. Ecosyst. Health Manag. 21, 229–233 (2018).

14. Munawar, M., Norwood, W., McCarthy, L. & Mayfield, C. In situ bioassessment of dredging and disposal activities in a contaminated ecosystem: Toronto Harbour. (1989) doi:10.1007/978-94-009-1896-2_62.

15. Dahmer, S. C., Matos, L. & Jarvie, S. Assessment of the degradation of aesthetics Beneficial Use Impairment in the Toronto and region Area of Concern. Aquat. Ecosyst. Health Manag. 21, 276–284 (2018).

16. Metro Toronto and Region Remedial Action Plan. Within Reach: 2015 Toronto an Region Remedial Action Plan Progress Report. (2016).

17. Burniston, D. & Waltho, J. Report on Sediment Quality in the Toronto Inner Harbour 2007. (2011).

18. Elbrecht, V., Vamos, E. E., Meissner, K., Aroviita, J. & Leese, F. Assessing strengths and weaknesses of DNA metabarcoding-based macroinvertebrate identification for routine stream monitoring. Methods Ecol. Evol. 8, 1265–1275 (2017).

19. Emilson, C. E. et al. DNA metabarcoding and morphological macroinvertebrate metrics reveal the same changes in boreal watersheds across an environmental gradient. Sci. Rep. 7, 12777 (2017).

20. Aylagas, E., Borja, Á., Muxika, I. & Rodríguez-Ezpeleta, N. Adapting metabarcoding-based benthic biomonitoring into routine marine ecological status assessment networks. Ecol. Indic. 95, 194–202 (2018).

21. Bush, A. et al. Studying ecosystems with DNA metabarcoding: lessons from biomonitoring of aquatic macroinvertebrates. Front. Ecol. Evol. 7, 434 (2019).

22. Serrana, J. M., Miyake, Y., Gamboa, M. & Watanabe, K. Comparison of DNA metabarcoding and morphological identification for stream macroinvertebrate biodiversity assessment and monitoring. Ecol. Indic. (2019).

23. Fernández, S., Rodríguez-Martínez, S., Martínez, J. L., Garcia-Vazquez, E. & Ardura, A. How can eDNA contribute in riverine macroinvertebrate assessment? A metabarcoding approach in the Nalón River (Asturias, Northern Spain). Environ. DNA 1, 385–401 (2019).

24. Hajibabaei, M. et al. Watered-down biodiversity? A comparison of metabarcoding results from DNA extracted from matched water and bulk tissue biomonitoring samples. PLOS ONE 14, e0225409 (2019).

25. Baird, D. J. & Hajibabaei, M. Biomonitoring 2.0: a new paradigm in ecosystem assessment made possible by next-generation DNA sequencing. Mol. Ecol. 21, 2039–2044 (2012).

26. Hajibabaei, M., Baird, D. J., Fahner, N. A., Beiko, R. & Golding, G. B. A new way to contemplate Darwin’s tangled bank: how DNA barcodes are reconnecting biodiversity science and biomonitoring. Philos. Trans. R. Soc. B Biol. Sci. 371, 20150330 (2016).

27. Beermann, A. J., Zizka, V. M. A., Elbrecht, V., Baranov, V. & Leese, F. DNA metabarcoding reveals the complex and hidden responses of chironomids to multiple stressors. Environ. Sci. Eur. 30, 26 (2018).

28. Bush, A. et al. DNA metabarcoding reveals metacommunity dynamics in a threatened boreal wetland wilderness. Proc. Natl. Acad. Sci. 117, 8539–8545 (2020).

29. Compson, Z. G. et al. Chapter two - Linking DNA metabarcoding and text mining to create network-based biomonitoring tools: a case study on boreal wetland macroinvertebrate communities. in Advances in Ecological Research (eds. Bohan, D. A., Dumbrell, A. J., Woodward, G. & Jackson, M.) vol. 59 33–74 (Academic Press, 2018).

30. Fernandes, K. et al. DNA metabarcoding—a new approach to fauna monitoring in mine site restoration. Restor. Ecol. 26, 1098–1107 (2018).

31. Fernandes, K. et al. Invertebrate DNA metabarcoding reveals changes in communities across mine site restoration chronosequences. Restor. Ecol. 27, 1177–1186 (2019).

32. Poikane, S. et al. Benthic macroinvertebrates in lake ecological assessment: A review of methods, intercalibration and practical recommendations. Sci. Total Environ. 543, 123–134 (2016).

33. Macher, J.-N. et al. Comparison of environmental DNA and bulk-sample metabarcoding using highly degenerate cytochrome c oxidase I primers. Mol. Ecol. Resour. 18, 1456–1468 (2018).

34. Marshall, N. T. & Stepien, C. A. Macroinvertebrate community diversity and habitat quality relationships along a large river from targeted eDNA metabarcode assays. Environ. DNA 2, 572–586 (2020).

35. Metro Toronto and Region Remedial Action Plan. Updates on Actions 2013-2014. (2013).

36. López-López, E. & Sedeño-Díaz, J. E. Biological indicators of water quality: The role of fish and macroinvertebrates as indicators of water quality. in Environmental Indicators (eds. Armon, R. H. & Hänninen, O.) 643–661 (Springer Netherlands, 2015). doi:10.1007/978-94-017-9499-2_37.

37. Berry, O. et al. A comparison of morphological and DNA metabarcoding analysis of diets in exploited marine fishes. (2015).

38. Sweeney, B. W., Battle, J. M., Jackson, J. K. & Dapkey, T. Can DNA barcodes of stream macroinvertebrates improve descriptions of community structure and water quality? J. North Am. Benthol. Soc. 30, 195–216 (2011).

39. Banerji, A. et al. Spatial and temporal dynamics of a freshwater eukaryotic plankton community revealed via 18S rRNA gene metabarcoding. Hydrobiologia 818, 71–86 (2018).

40. Porter, T. M. et al. Widespread occurrence and phylogenetic placement of a soil clone group adds a prominent new branch to the fungal tree of life. Mol. Phylogenet. Evol. 46, 635–644 (2008).

41. Rosling, A. et al. Archaeorhizomycetes: unearthing an ancient class of ubiquitous soil fungi. Science 333, 876–879 (2011).

42. Mandaville, S. M. Benthic Macroinvertebrates in Freshwaters-Taxa Tolerance Values, Metrics, and Protocols. 128 http://lakes.chebucto.org/H-1/tolerance.pdf (2002).

43. Trzcinski, M. K. et al. The effects of food web structure on ecosystem function exceeds those of precipitation. J. Anim. Ecol. 85, 1147–1160 (2016).

44. Liu, X. & Wang, H. Contrasting patterns and drivers in taxonomic versus functional diversity, and community assembly of aquatic plants in subtropical lakes. Biodivers. Conserv. (2018).

45. Kovalenko, K. E., Brady, V. J., Ciborowski, J. J. H., Ilyushkin, S. & Johnson, L. B. Functional changes in littoral macroinvertebrate communities in response to watershed-level anthropogenic stress. PLOS ONE 9, e101499 (2014).

46. Luiza-Andrade, A., Montag, L. F. de A. & Juen, L. Functional diversity in studies of aquatic macroinvertebrates community. Scientometrics 111, 1643–1656 (2017).

47. MacMillan, G. A., Chételat, J., Heath, J. P., Mickpegak, R. & Amyot, M. Rare earth elements in freshwater, marine, and terrestrial ecosystems in the eastern Canadian Arctic. Environ. Sci. Process. Impacts 19, 1336–1345 (2017).

48. Pastorino, P. et al. Macrobenthic invertebrates as tracers of rare earth elements in freshwater watercourses. Sci. Total Environ. 698, 134282 (2020).

49. Kulaš, A. et al. Ciliates (Alveolata, Ciliophora) as bioindicators of environmental pressure: A karstic river case. Ecol. Indic. 124, 107430 (2021).

50. Persaud, D., Lomas, T., Boyd, D. & Mathai, S. Historical development and quality of the Toronto waterfront sediments. (1985).

51. Milani, D. & Grapentine, L. Assessment of sediment quality in the Bay of Quinte Area Of Concern. (2000).

52. Reynoldson, T. B., Bailey, R. C., Day, K. E. & Norris, R. H. Biological guidelines for freshwater sediment based on BEnthic Assessment of SedimenT (the BEAST) using a multivariate approach for predicting biological state. Aust. J. Ecol. (1995).

53. Geller, J., Meyer, C., Parker, M. & Hawk, H. Redesign of PCR primers for mitochondrial cytochrome c oxidase subunit I for marine invertebrates and application in all-taxa biotic surveys. Mol. Ecol. Resour. 13, 851–861 (2013).

54. Leray, M. et al. A new versatile primer set targeting a short fragment of the mitochondrial COI region for metabarcoding metazoan diversity: application for characterizing coral reef fish gut contents. Front. Zool. 10, 34 (2013).

55. Zhan, A. et al. High sensitivity of 454 pyrosequencing for detection of rare species in aquatic communities. Methods Ecol. Evol. 4, 558–565 (2013).

56. Gibson, J. et al. Simultaneous assessment of the macrobiome and microbiome in a bulk sample of tropical arthropods through DNA metasystematics. Proc. Natl. Acad. Sci. 111, 8007–8012 (2014).

57. Gibson, J. F. et al. Large-scale biomonitoring of remote and threatened ecosystems via high-throughput sequencing. PLOS ONE 10, e0138432 (2015).

58. Porter, T. M. & Hajibabaei, M. METAWORKS: A flexible, scalable bioinformatic pipeline for multi-marker biodiversity assessments. bioRxiv 2020.07.14.202960 (2020).

59. Köster, J. & Rahmann, S. Snakemake—a scalable bioinformatics workflow engine. Bioinformatics 28, 2520–2522 (2012).

60. Anon. Conda. (2016).

61. Porter, T. M. & Hajibabaei, M. Automated high throughput animal CO1 metabarcode classification. Sci. Rep. 8, 4226 (2018).

62. Porter, T. M. Eukaryote CO1 Reference Set For The RDP Classifier. (Zenodo, 2017). doi:10.5281/zenodo.4741447.

63. Porter, T. M. SILVA 18S Reference Set For The RDP Classifier. (Zenodo, 2018). doi:10.5281/zenodo.4741433.

64. R Core Team. R: A language and environment for statistical computing. (2020).

65. Wickham, H. ggplot2: Elegant Graphics for Data Analysis. (Springer-Verlag, 2009). doi: 10.1007/978-0-387-98141-3.

66. Oksanen, J. et al. vegan: Community Ecology Package. (2020).

67. Komsta, L. & Novomestky, F. moments: Moments, cumulants, skewness, kurtosis and related tests. (2015).

68. Harrell, F. & functions), C. D. (contributed several functions and maintains latex. Hmisc. (2021).

69. U.S. Environmental Protection Agency. Freshwater Biological Traits Database. (2012).

70. Schmidt-Kloiber, A. & Hering, D. www.freshwaterecology.info – An online tool that unifies, standardises and codifies more than 20,000 European freshwater organisms and their ecological preferences. Ecol. Indic. 53, 271–282 (2015).

71. Moog, O. Fauna Aquatica Austriaca - Catalogue for autecological classification of Austrian aquatic organisms. (1995).

72. Usseglio-Polatera, P., Tachet, H., Richoux, P. & Bournaud, M. Invertébrés d’eau douce - systématique, biologie, écologie. (2000).

73. Nally, R. M. & Walsh, C. J. Hierarchical Partitioning Public-domain Software. Biodivers. Conserv. (2004) doi:10.1023/B:BIOC.0000009515.11717.0b.

