## Supplementary information for "Multi-marker DNA metabarcoding detects suites of environmental gradients from an urban harbour"

Supplementary Information Text

**Extended Technical Description of Sampling Methods**

**Methods**

*Surface and overlying water sampling*

Prior to sediment collection, water samples were obtained using a 1 litre van Dorn sampler, taken at 0.5 meters below the surface and 0.5 meters from the bottom. For this study, we only focused on water collected 0.5 meters from the bottom. Water was dispensed directly into two separate bottles; for major ions (500 mL polyethylene bottle) and nutrients (120 mL glass). Physical data (temperature, conductivity, pH and dissolved oxygen) was collected using water collected at 0.5 m below the water surface and at 0.5 m from the bottom ^1–3^.

Major ions in water were performed by the National Laboratory for Environmental Testing (NLET; Burlington, ON). For calcium, magnesium, sodium and potassium, a sample aliquot was combined with an ionization suppressant and releasing agent solution and aspirated into the air-acetylene flame of an atomic absorption spectrophotometer ^4^.

For the analysis of fluoride, chloride, nitrate, and sulphate, the water sample was injected into the eluent stream, which was pumped through two columns packed with low-capacity anion exchange resin in the form of CO_3_/HCO_3_. The ions of interest (chloride, nitrate and sulphate) separated based on their affinity for converting the anions in the sample to their respective acid forms (e.g., HCl, HNO_3_). The concentrations of these separated anions were determined by measuring their respective conductivities using a conductivity detector. Anions are identified by their retention times compared to the known standards ^4,5^.

*Sediment sampling*

Sediment samples which were analysed for total organic carbon (TOC), total inorganic carbon (TIC) and loss on ignition (LOI), were freeze dried prior to analysis. Freeze drying, TOC, TIC, LOI and grain size distribution were conducted by Natural Resources Canada (Terrain Sciences Division Ottawa, ON). TOC was analyzed by Leco-Cr-412 and grain size fractions were determined using a Lecotrac Particle Size Analyzer LT100. In addition, trace metals analysis was performed by the NLET (Burlington, ON). Trace metal analysis were determined by a hot aqua-regia extraction with measurement by ICP-AES. Total mercury was determined by digestion with hot nitric acid and hydrochloric acid followed with measurement by cold vapour atomic adsorption spectrometer ^6^. In addition, presence of PCB aroclors, total PCBs and PCB congeners, chlorinated pesticides, chlorobenzenes, technical toxaphene and toxaphene congeners/parlars was determined by AXYS Technologies (AXYS, Sidney, British Columbia, Canada).

The overall sediment quality results were compared to the Canadian Environmental Quality Guidelines ^6^. The CCME sediment quality guidelines provide scientific benchmarks, for evaluating the potential for observing adverse biological effects in aquatic systems ^6^. The guidelines are derived from available toxicological information. A lower value, referred to as the threshold effect level (TEL), represents the concentration below which adverse biological effects are expected to occur rarely. The upper value, referred to as the probable effect level (PEL), represents the level above which adverse effects are expected to occur frequently. Fewer than 25% of adverse effects occur below the TEL, and more than 50% of adverse effects occur above the PEL ^7^.

*Benthos samples*

Benthos samples were collected for both morphology and metabarcoding analyses. To achieve this, a 40 cm x 40 cm mini-box core was lowered twice at each station to obtain sediment and benthos samples. From one box core at each station, subsamples were collected in triplicate using a sampling technique designed to minimize DNA contamination (i.e., new sterile gloves, spatula, and mini-Whirl-Pak® for each sample). A disposable spatula was used to collect each benthos sample (5-10 mL of sediment per sample) from the box corer and placed into a Whirl-Pak®. DNA samples were stored on dry ice on the boat and placed in a -80°C freezer upon return to Canada Centre for Inland Waters (CCIW).

Twice during the 3-day sampling period, a box-corer field blank was taken by pouring MilliQ® water along the inside of box corer and collecting the respective water in a 15 mL Falcon tube for DNA analysis. Similarly, hose blanks were also taken twice during the 3-day sampling period by pouring MilliQ® water on the hose nozzle and collecting the water in a 15 mL Falcon tube for DNA analysis. Hose blanks were taken after the hose was rinsed with site water, as per normal practice between sites.

For the second box corer at each station, benthos community sub samples were collected from the box core with five 10 cm (6.5 cm diameter) acrylic tubes and each dispensed into plastic containers with 5% formalin for preservation on morphological characters. Samples were agitated thoroughly to ensure formalin saturated the sediment. In the laboratory, samples were sieved through a 250 µm mesh ^3-5^. The formalin was decanted and replaced with 70% ethanol. The top 3 cm of remaining sediment in the box core was skimmed off the top using a stainless-steel spoon, passed through a 250 µm mesh screen into a glass bowl and stirred for 2 minutes to further homogenate the sample for downstream sediment analysis ^7^.

*METAWORKS bioinformatic processing*

SeqPrep v1.3.2 ^8^ was used to pair raw reads requiring a minimum Phred score of 20 to ensure 99% base-calling accuracy and a minimum of 25 bp overlap. CUTADAPT v2.6 was used to trim primers from sequences, using a minimum Phred score of 20 at the ends, leaving a minimum fragment length of at least 150 base pairs, no more than 3 Ns permitted ^9^. Global exact sequence variant (ESV) ^10^ analysis was performed on the primer-trimmed reads. Reads were dereplicated using the ‘derep_fulllength’ command with the ‘sizein’ and ‘sizeout’ options of VSEARCH v2.14.1 ^11^. VSEARCH was also used to denoise the data using the unoise3 algorithm ^12^. These steps were taken to correct sequences with errors and remove rare reads (singletons or doubletons) ^13^. Putative chimeric sequences were removed using the ‘uchime3_denovo’ algorithm in VSEARCH. An ESV x sample table was created using the ‘search_exact’ method in VSEARCH.

COI ESVs were classified using the COI Classifier (v4) available from <https://github.com/terrimporter/CO1Classifier/releases/tag/v4>, comprised of a curated reference sequence set mined from BOLD ^14^ and GenBank ^15^. 18S ESVs were classified using the 18S Eukaryota Classifier (v4.1) available from <https://github.com/terrimporter/18SClassifier/releases/tag/v4.1>, created from the SILVA 138 SSU Ref Nr99 release ^16^. Sequences without a genus rank assignment from SILVA were excluded from the reference database. Both COI and 18S classifiers were used with the RDP classifier v2.12, using a naive Bayesian algorithm. For COI, we used a 0.20 bootstrap support cutoff at the family rank (99% correct assignments expected assuming the query sequences are represented in the reference database), a 0.30 cutoff at the genus rank (99% correct expected) and a 0.70 support cutoff at the species rank (95% correct expected) ^17^. For 18S, we used a 0.00 bootstrap support cutoff at the phylum to class rank (80% correct expected for order, 90% correct expected for class and phylum, 95% correct for kingdom, 99% correct for domain), and a 0.20 cutoff at family rank (80% correct expected), 0.70 cutoff at genus rank (80% correct expected).

For COI, putative pseudogenes were identified and removed in the METAWORKS pipeline as follows: Denoised ESVs were translated into every possible reading frame on the plus strand using ORFfinder v0.4.3 ^18^ keeping the longest open reading frame (ORF). Amino acid ORFs were used for hidden Markov model (HMM) profile analysis. ORFs with a full sequence bit score lower than the 25th percentile - 1.5 * interquartile length were excluded as putative pseudogenes or sequences with errors that cause internal stop codons.

**Figure S1: COI metabarcoding can provide finer scale resolution and more complete sampling from bulk benthos DNA samples.** We compared DNA metabarcoding and traditional benthic sampling by showing a) median taxonomic assignment resolution and b) sampling effort.

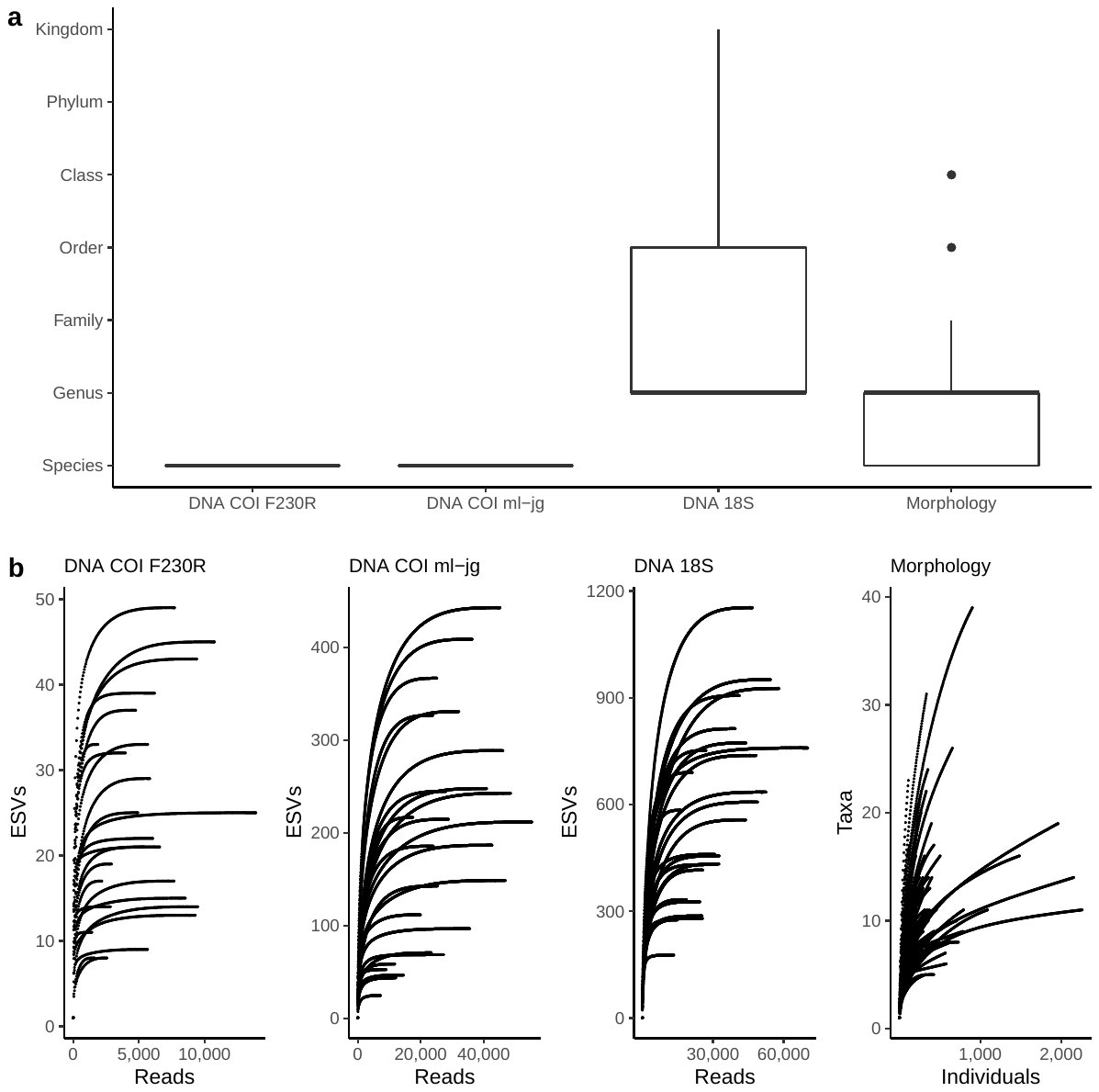

**Figure S2: Correlation between water physical-chemical features and sediment contaminants.** Pearson correlations are shown if greater than 0.70 with a p-value < 0.05.

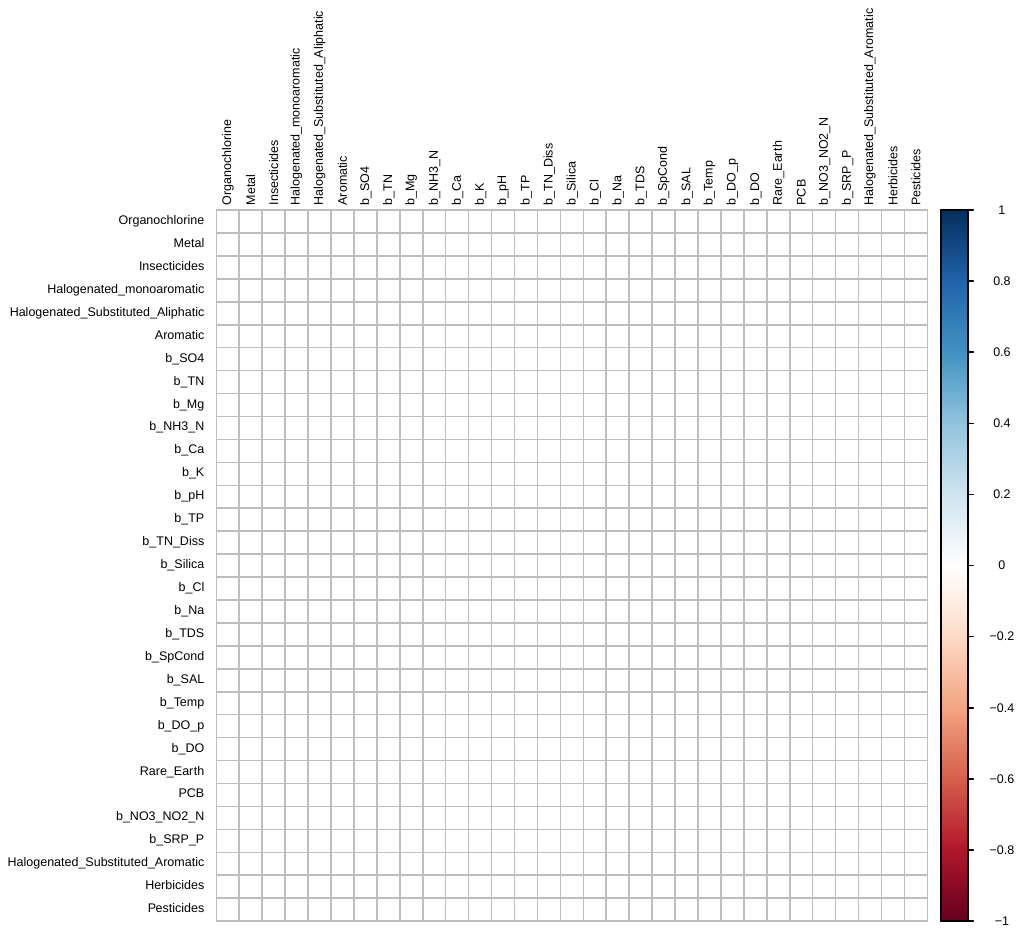

Table S1. GPS coordinates for each station, detailing location of each station within Toronto Harbour.

| **Station** | **Location** | **Latitude** | **Longitude** |
| --- | --- | --- | --- |
| 1346 | Bathurst Quay | 43.6356 | -79.3959 |
| 1362 | Inner Harbour Centre | 43.6383 | -79.3754 |
| 1363 | Inner Harbour Centre | 43.6376 | -79.3726 |
| 1364 | Inner Harbour Centre | 43.6354 | -79.3712 |
| 1365 | Inner Harbour Centre | 43.6325 | -79.3701 |
| 1366 | Inner Harbour Centre | 43.6289 | -79.367 |
| 1371 | Inner Harbour East | 43.6392 | -79.359 |
| 1372 | Inner Harbour East | 43.6362 | -79.3527 |
| 1380 | Inner Harbour East | 43.6407 | -79.3559 |
| 6699 | Inner Harbour East | 43.6392 | -79.3626 |
| 1012 | Inner Harbour West | 43.634 | -79.3956 |
| 1349 | Inner Harbour West | 43.6358 | -79.3915 |
| 1355 | Inner Harbour West | 43.6368 | -79.384 |
| 1358 | Inner Harbour West | 43.6303 | -79.3809 |
| 1359 | Inner Harbour West | 43.6261 | -79.379 |
| 1761 | Inner Harbour West | 43.6297 | -79.3873 |
| 1765A | Inner Harbour West | 43.627 | -79.3831 |
| 1772 | Inner Harbour West | 43.6234 | -79.3775 |
| 1375 | Keating Channel | 43.6442 | -79.3598 |
| 1379A | Keating Channel | 43.6467 | -79.3554 |
| 1370 | Queen Elizabeth Docks | 43.6412 | -79.3623 |
| 1381 | Ship Channel | 43.6442 | -79.344 |
| 1384 | Ship Turning Basin | 43.6523 | -79.3342 |
| 1401 | Toronto Island | 43.6219 | -79.3879 |
| 1403 | Toronto Island | 43.6201 | -79.3812 |

Table S2. Summary of read and ESV counts for each primer set and in total.

|  | **Counts** | **COI - F230R** | **COI - ml-jg** | **18S** | **Total** |
| --- | --- | --- | --- | --- | --- |
| **Raw paired-end reads** | 18,790,832 x 2 |  |  |  | 18,790,832 x 2 |
| **Paired reads** | 17,240,306 |  |  |  | 17,240,306 |
| **Primer trimmed sequences** |  | 3,057,498 | 7,331,237 | 4,232,625 | 14,621,360 |
| **ESVs (reads)** |  | 182 (296,798) | 2,544 (1,312,427) | 6,603 (1,285,245) | 9,329 (2,894,470) |

**Table S3.** A checklist of macroinvertebrate families detected using conventional methods and metabarcoding in Toronto Harbour. See separate Excel sheet.

**Table S4.** 2018 Toronto Harbour environmental characteristics. Sediment contaminant values in red exceed the Canadian Federal Probable Effect Level (PEL).

| **Nutrients/major ions measured from water near the bottom of the harbour** | **Mean** | **Minimum** | **Maximum** |  |
| --- | --- | --- | --- | --- |
| Temperature (ºC) | 15.82 | 0 | 17.1 |  |
| Dissolved oxygen (%) | 90.744 | 80.1 | 98.6 |  |
| Dissolved oxygen (mg/L) | 8.8596 | 7.76 | 9.89 |  |
| Specific conductance (µS/cm) | 338.382 | 299.5 | 429 |  |
| Total dissolved solids (g/L) | 0.22 | 0.2 | 0.28 |  |
| Salinity (ppt) | 0.1616 | 0.14 | 0.21 |  |
| pH | 8.0472 | 7.76 | 8.29 |  |
| CaCO3 (mg/L) | 94.584 | 91.6 | 112 |  |
| Ammonia (mg/L) | 0.07496 | 0.025 | 364 |  |
| Calcium (mg/L) | 36.424 | 34 | 47.3 |  |
| Chloride (mg/L) | 34.812 | 24.3 | 90.9 |  |
| Fluoride (mg/L) | 0.1112 | 0.11 | 0.12 |  |
| Magnesium (mg/L) | 9.0448 | 8.66 | 9.12 |  |
| Nitrate/Nitrite (mg/L) | 0.30132 | 0.024 | 0.844 |  |
| Total nitrogen (mg/L) | 0.29576 | 0.223 | 0.75 |  |
| Total nitrogen dissolved (mg/L) | 0.29972 | 0.212 | 0.723 |  |
| Phosphorus SRP (mg/L) | 0.010752 | 0.0011 | 0.0271 |  |
| Total phosphorus (mg/L) | 0.02814 | 0.0117 | 0.133 |  |
| Potassium (mg/L) | 1.86 | 1.7 | 2.56 |  |
| Silica (mg/L) | 1.0948 | 0.51 | 3.72 |  |
| Sodium (mg/L) | 21.276 | 14.8 | 55.9 |  |
| Sulphate (mg/L) | 24.068 | 23.2 | 25.8 |  |
| **Sediment contaminants** |  |  |  | **PEL** |
| Aromatic hydrocarbons (ng/g) | 13083.11 | 2498 | 98127 | 7106.9 |
| Halogenated (substituted aliphatic) hydrocarbons (ng/g dwt) | 0.0648 | 0.02 | 0.25 |  |
| Halogenated (substituted aromatic) hydrocarbons (ng/g dwt) | 0.5596 | 0.19 | 1.38 |  |
| Halogenated monoaromatic hydrocarbons (ng/g dwt) | 3.2032 | 0.8 | 9.85 |  |
| Herbicides (ng/g dwt) | 0.6208 | 0.11 | 1.66 |  |
| Insecticides | 35.5464 | 10.06 | 176.27 |  |
| DDT | 3.92 | 1.69 | 17 | 4.77 |
| DDE | 7.45 | 2.1 | 21.1 | 6.75 |
| Metals (µg/g) | 63129.68 | 22081.11 | 89070.27 |  |
| Arsenic | 6.32 | 2 | 10 | 17 |
| Cadmium | 1.28 | 3 | 3.9 | 3.5 |
| Chromium | 68.28 | 17 | 135 | 90 |
| Copper | 92.68 | 23 | 194 | 97 |
| Lead | 102.18 | 15.8 | 381 | 91.3 |
| Manganese | 715.6 | 276 | 1480 |  |
| Mercury | 0.26 | 0.04 | 0.97 | 0.486 |
| Nickel | 28.69 | 8.2 | 45.9 |  |
| Zinc | 305 | 66 | 583 | 315 |
| Metalloids (µg/g) | 0.02 | 0.02 | 0.02 |  |
| Organochlorines (ng/g dwt) | 9.0768 | 1.81 | 69.03 |  |
| Hexachlorocyclohexane | 3.3 | 0.53 | 11.2 | 1.38 |
| PCBs (ng/g dwt) | 152.232 | 34.68 | 537.65 | 277 |
| Pesticides (ng/g dwt) | 28.1324 | 8 | 66.17 |  |
| DDD | 8.49 | 2.1 | 32.6 | 8.51 |
| Rare earth metals (µg/g) | 69.0992 | 31.18 | 87.1 |  |
| Cerium | 47.53 | 21.6 | 60.3 |  |
| Lanthanum | 21.57 | 9.58 | 26.8 |  |

**Table S5. Water physical-chemical features that make significant independent contributions (p-value < 0.05) partitioning variance.**

| **Predictors** | **Individual contribution (%)** | **Individual / Joint contribution *** | **Metric type **** | **Metric responses ***** | **Sampling method** |
| --- | --- | --- | --- | --- | --- |
| Temperature | 61.9 | 7.6 | D | mi.richness.dna | COI |
| Ammonia | 56.3 | 2.2 | D | Bosminidae.dna | COI |
| Ammonia | 56.0 | 1.7 | D | Naididae.dna | COI |
| Ammonia | 50.4 | 3.0 | D | Candonidae.dna | COI |
| Temperature | 49.6 | 3.1 | D | Haptoria.dna | 18S |
| Ammonia | 49.2 | 3.3 | D | Thoracosphaeraceae.dna | 18S |
| Ammonia | 47.3 | 5.7 | D | Hydridae.morph | Morphology |
| Total Nitrogen | 38.2 | -8.9 | D | Limnesiidae.morph | Morphology |
| pH | 23.6 | -2.0 | D | Limnesiidae.morph | Morphology |
| Ammonia | 56.9 | 24.1 | F | PA.morph | Morphology |
| Sulphate | 32.8 | 1.1 | F | PR.dna | COI |
| Temperature | 32.2 | -44.7 | F | HB.morph | Morphology |
| Sulphate | 31.7 | 2.0 | F | CF.dna | COI |
| Ammonia | 29.7 | -2.8 | F | SH.morph | Morphology |
| Temperature | 29.2 | 12.4 | F | CG.morph | Morphology |
| Ammonia | 29.0 | -2.3 | F | Other.morph | Morphology |
| Ammonia | 28.9 | -3.2 | F | CG.morph | Morphology |
| Temperature | 27.6 | 29.5 | F | SH.morph | Morphology |
| Temperature | 23.2 | -460.9 | F | Other.morph | Morphology |

* Negative joint effects indicate that this predictor acts as a suppressor on other variables

** Metric type: diversity (D), functional (F)

*** macroinvertebrate (mi), conventional morphology (morph), DNA metabarcoding (dna), parasites (PA), predators (PR), herbivores (HB), collector-filterers (CF), shredders (SH), collector-gatherers (CG)

**Table S6. Sediment contaminants that make significant independent contributions (p-value < 0.05) partitioning variance.**

| **Predictors*** | **Individual contribution (%)** | **Individual / Joint contribution **** | **Metric type ***** | **Metric responses ****** | **Sampling method** |
| --- | --- | --- | --- | --- | --- |
| Aromatic | 62.7 | -2.1 | D | Candonidae.dna | COI |
| Metals | 61.6 | 2.9 | D | mi.richness.dna | COI |
| Aromatic | 57.1 | 2.5 | D | Cyprididae.dna | COI |
| Metals | 55.1 | 13.1 | D | Thoracosphaeraceae.dna | 18S |
| Aromatic | 44.6 | 1.6 | D | Limnesiidae.morph | Morphology |
| Metals | 41.4 | 4.9 | D | euk.richness.dna | 18S |
| Aromatic | 41.1 | 2.0 | D | Naididae.morph | Morphology |
| Orgchl | 36.9 | 3.0 | D | euk.richness.dna | 18S |
| Aromatic | 36.7 | -1.4 | D | Bosminidae.dna | COI |
| Metals | 33.8 | -12.4 | D | Oligohymenophorea.dna | 18S |
| Insecticides | 32.2 | 1.3 | D | Limnesiidae.morph | Morphology |
| Metals | 30.9 | 2.0 | D | Naididae.morph | Morphology |
| Insecticides | 24.1 | -1.0 | D | Candonidae.dna | COI |
| Orgchl | 21.8 | -3.0 | D | Bosminidae.dna | COI |
| Metals | 18.0 | -3.1 | D | Bosminidae.dna | COI |
| Orgchl | 59.2 | 6.6 | F | PA.dna | COI |
| Orgchl | 46.2 | 5.5 | F | CF.dna | COI |

* Aromatic hydrocarbons (Aromatic), organochlorines (Orgchl)

** Negative values indicate this predictor acts as a suppressor on other variables

*** Metric type: diversity (D), functional (F)

**** macroinvertebrate (mi), conventional morphology (morph), DNA metabarcoding (dna), eukaryote (euk), parasites (PA), collector-filterers (CF)

**SI References**

1.	Reynoldson, T. B., Bailey, R. C., Day, K. E. & Norris, R. H. Biological guidelines for freshwater sediment based on BEnthic Assessment of SedimenT (the BEAST) using a multivariate approach for predicting biological state. *Aust. J. Ecol.* (1995).

2.	Milani, D. & Grapentine, L. “Assessment of sediment quality in the Bay of Quinte Area Of Concern” (Environment Canada, 2000).

3.	Burniston, D. & Waltho, J. “Report on Sediment Quality in the Toronto Inner Harbour 2007” (Environment Canada, 2011).

4.	National Laboratory for Environmental Testing (NLET), “Manual of Analytical Methods, Major Ions and Nutrients” (Environment Canada, 1997).

5.	Dove, A. Long-term trends in major ions and nutrients in Lake Ontario. *Aquat. Ecosyst. Health Manag.* **12**, 281–295 (2009).

6.	Canadian Council of Ministers of the Environment, CCME Summary Table (2001) (August 24, 2020).

7.	Marvin, C.H. *et al.* Surficial Sediment Contamination in Lakes Erie and Ontario: A Comparative Analysis. *J. Gt. Lakes Res.* **28**, 437–450 (2002).

8.	St. John, J. SeqPrep. *Retrieved https://github.com/jstjohn/SeqPrepreleases* (2016).

9.	Martin, M. Cutadapt removes adapter sequences from high-throughput sequencing reads. *EMBnet.journal* **17**, 10–12 (2011).

10.	Callahan, B. J., McMurdie, P. J. & Holmes, S. P. Exact sequence variants should replace operational taxonomic units in marker-gene data analysis. *ISME J.* **11**, 2639–2643 (2017).

11.	Rognes, T., Flouri, T., Nichols, B., Quince, C. & Mahé, F. VSEARCH: A versatile open-source tool for metagenomics. *PeerJ* **2016**, 1–22 (2016).

12.	Edgar R. C., UNOISE2: improved error-correction for Illumina 16S and ITS amplicon sequencing. *bioRxiv* (2016) https:/doi.org/10.1101/081257.

13.	Reeder, J. & Knight, R. The “rare biosphere”: A reality check. *Nat. Methods* **6**, 636–637 (2009).

14.	Ratnasingham, S. & Hebert, P. D. N. A DNA-Based Registry for All Animal Species: The Barcode Index Number (BIN) System. *PLOS ONE* **8**, e66213 (2013).

15.	Benson, D. A. *et al.* GenBank. *Nucleic Acids Res.* **41**, D36–D42 (2013).

16.	Pruesse, E. *et al.* SILVA: a comprehensive online resource for quality checked and aligned ribosomal RNA sequence data compatible with ARB. *Nucleic Acids Res.* **35**, 7188–7196 (2007).

17.	Porter, T. M. & Hajibabaei, M. Automated high throughput animal CO1 metabarcode classification. *Sci. Rep.* **8**, 4226 (2018).

18.	National Center for Biotechnology Information, *Open Reading Frame Finder (RRID:SCR_016643). Software tool to search for open reading frames (ORFs) in the DNA sequence.* (2020).
